# EFFECTS OF AQUEOUS EXTRACT OF *CYMBOPOGON CITRATUS* ON BLOOD GLUCOSE, BODY WEIGHT, AND PANCREATIC ANTIOXIDANT ACTIVITIES IN NORMAL AND STREPTOZOTOCIN-INDUCED DIABETIC RATS

**DOI:** 10.1101/2025.10.24.684396

**Authors:** Lincoln Osas Afere, Success Oseze Iziduh

**Affiliations:** Department of Biochemistry, Faculty of Life Sciences, University of Benin, Benin City, Nigeria

**Author notes:** Corresponding Author: Dr. Lincoln Osas Afere.

**Keywords:** *Cymbopogon Citratus*, Antioxidants, Diabetes Mellitus, Streptozotocin, Blood Glucose

## Abstract

*Cymbopogon citratus* (lemongrass) is widely used in folk medicine as an antioxidant and for pancreas ailments. This study investigated the effects of its aqueous extract on diabetic and oxidative status in normal and streptozotocin-induced (STZ-induced) diabetic rats. Diabetes was induced by a single intraperitoneal injection of STZ (45 mg/kg body weight) in the diabetic groups. Treated groups received oral administration of aqueous lemongrass extract at a dose of 400 mg/kg body weight daily for 21 days. Biochemical studies measured body weight, fasting blood glucose (FBG) levels, and pancreatic activities of superoxide dismutase (SOD) and catalase (CAT). Relative to the normal control (NC) rats, the extract caused a significant decrease (*P <* 0.05) in final blood glucose in normal treated (NT) rats. STZ injection resulted in a significant increase (*P <* 0.05) in FBG in diabetic control (DC) rats compared to NC rats. However, FBG in diabetic treated (DT) rats was not statistically different (*P >* 0.05) compared to DC rats. The extract caused an insignificant decrease (*P >* 0.05) in body weight in NT rats compared to NC rats, while STZ caused a significant decrease (*P <* 0.05) in DC rats compared to NC rats. Weight gained by DT rats was not statistically different (*P >* 0.05) compared to DC rats. SOD and catalase activities in the pancreatic tissues of NT and DT rats were not statistically different (*P >* 0.05) when compared to NC and DC groups, as the activities of all four groups were statistically similar (*P >* 0.05). The aqueous lemongrass extract was ineffective against STZ-induced diabetes, as it did not significantly increase body weight nor decrease blood glucose levels in the diabetic rats. However, it did cause hypoglycemia in the normal rats. No oxidative damage to the pancreas was observed from the aqueous lemongrass extract.

## 1 Introduction

Diabetes mellitus is a group of diseases leading to high blood sugar levels (hyperglycemia). There are three main types: Type 1 (pancreas produces little to no insulin), Type 2 (body resists insulin or doesn’t synthesize enough), and Gestational diabetes (glucose intolerance during pregnancy) (Saucedo et al., 2023). In Type 2 diabetes, cells become resistant to insulin, causing glucose levels to rise in the bloodstream. Weight loss is a symptom of diabetes due to the excessive use of fat and proteins as substitutes for unusable glucose (Junior, 2024).

*Cymbopogon citratus*, commonly known as lemongrass, is a genus of grasses native to tropical Asia and southern India, often used in folk medicine for inflammation, indigestion, and as an antioxidant. The active components in its essential oil include citral, myrcene, limonene, and geraniol (Aluyor and Oboh, 2014).

Streptozotocin (STZ) is an antibiotic widely used experimentally to induce a model of Type 1 diabetes (TIDM) by causing pancreatic islet *β*-cell destruction (Abdollahi and Hosseini, 2014). STZ is transported into the *β*-cell via glucose transporter 2 (GLUT2), leading to DNA alkylation, depletion of NAD^+^ and ATP, and the production of toxic superoxide, hydroxyl, and nitric oxide radicals, ultimately causing *β*-cell necrosis (Szkudelski, 2001).

Superoxide Dismutase (SOD) and Catalase (CAT) are antioxidant enzymes. SOD catalyzes the dismutation of superoxide radicals 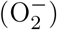 into molecular oxygen (O_2_) and hydrogen peroxide (H_2_O_2_), while CAT is found in aerobic organisms and catalyzes H_2_O_2_ into water and oxygen (Chen et al., 2023; Zheng et al., 2023).

The objective of this study was to determine the effects of an aqueous extract of *Cymbopogon citratus* on normal and STZ-induced diabetic rats by evaluating changes in blood glucose, body weight, and pancreatic SOD and CAT activities.

## 2 Materials and Methods

### 2.1 Materials

- **Plant:** *Cymbopogon citratus* was purchased from Uselu Market, Benin City, and authenticated (voucher number: UBH-C451).
- **Animals:** Forty-five (45) male albino Wistar rats with initial mean weights between 120–153 g were purchased from the Department of Anatomy, University of Benin. They were acclimatized for 7 days, fed grower pellets and water *ad libitum*.
- **Chemicals:** Streptozotocin (Sigma, London) and a Catalase and Superoxide Dismutase Assay Kits (Randox Laboratories Ltd, UK) were used.

### 2.2 Preparation of Aqueous Extract

One hundred grams (100 g) of dried, grounded lemongrass was boiled in 2 liters of distilled water for 10 minutes. The mixture was filtered using a muslin cloth, and the filtrate was dried using a rotary evaporator at 60°C to obtain the solid extract, which was stored at 4°C.

### 2.3 Induction of Diabetes and Treatment

STZ (45 mg) was dissolved in 1 ml of 0.1 M sodium citrate buffer (pH 4.5). Diabetes was induced by intraperitoneal injection of the STZ solution at a volume of 1ml/kg body weight. Rats were divided into four groups for a 21-day treatment period (Sadique et al., 1987):

- **Normal Control (NC, 5 rats)** - Rats were injected with the buffer only and weren’t treated with the aqueous extract.
- **Normal Treated (NT, 10 rats)** - Rats were injected with buffer and treated with the aqueous extract at a dose of 8 ml/kg b. wt. (see appendix I)
- **Diabetic Control (DC, 15 rats)** - Rats were injected with Streptozotocin/buffer solution at a dose of 1ml/kg b. wt. (see appendix I) but weren’t treated with the extract.
- **Diabetic Treated (DT, 15 rats)** - Rats were injected with Streptozotocin/buffer solution (at a dose of 1ml/kg b. wt.) and treated with the extract (at a dose of 8 ml/kg b. wt.)

After the injections, the rats were provided free access to 10% sucrose water for the next 24 hours. The blood glucose level of each rat was recorded every 7 days for the next 21 days.

### 2.4 Sample Collection and Biochemical Assays

Animals were sacrificed 24 hours after the last treatment, and the entire pancreas was excised and weighed.

- **Fasting Blood Glucose (FBG):** Measured every 7 days using a glucometer on blood obtained from the tail-tip after an overnight fast (Barham and Trinder, 1972).
- **Body Weight:** Initial and final weights were recorded using a weighing balance.
- **Pancreatic SOD and Catalase (CAT) Activity:** Determined using the pancreatic tissue homogenate supernatant, based on the methods described in the Randox kits (Misra and Fridovich, 1972; Cohen et al., 1970).

### 2.5 Statistical Analysis

Results are presented as mean *±* SEM. Data were analyzed using one-way Analysis of Variance (ANOVA) and Least Square Difference (LSD) post-hoc test (SPSS version 17). Statistical significance was set at *P <* 0.05.

## 3 Results

### 3.1 Body weight

Table 3.1 shows the initial, final and percentage change in weight of the corresponding groups within the 21 days treatment period. One of the symptoms of diabetes mellitus type 2 is weight loss, which is characterized by the excessive usage of fat and proteins as substitute for glucose which the body is unable to utilize. We observed promising levels of weight change in the diabetic treated groups as compared to the diabetic control groups.

**Table 1:** Change in body weight for the 21 days treatment period.

| GROUPS | Initial weight (g) | Final weight (g) | % wt. loss/gain |
| --- | --- | --- | --- |
| Normal control ($n = 5$) | $121.8 \pm 4.35^a$ | $189 \pm 5.62^b$ | +55.2% |
| Normal treated ($n = 5$) | $126.51 \pm 2.29^a$ | $185.2 \pm 6.98^b$ | +46.4% |
| Diabetic control ($n = 13$) | $141.15 \pm 1.68^c$ | $156.39 \pm 6.43^c$ | +10.8% |
| Diabetic treated ($n = 11$) | $148.97 \pm 2.89^c$ | $157.46 \pm 9.47^c$ | +5.7% |

Values are in mean ± SEM. Data with the same superscripts are not significantly different ( $p > 0.05$ ) while data with different superscripts are significantly different ( $p < 0.05$ ).

### 3.2 Fasting Blood Glucose

Table 3.2 shows the glucose levels of the corresponding groups throughout the 21 days treatment cycle. The basal fasting blood glucose of the rats in all the groups were not significantly different, but the animals in the diabetic groups were very responsive to the STZ treatment after 48hrs as there was a significant increase in the glucose levels as compared to the normal groups.

**Table 2:**
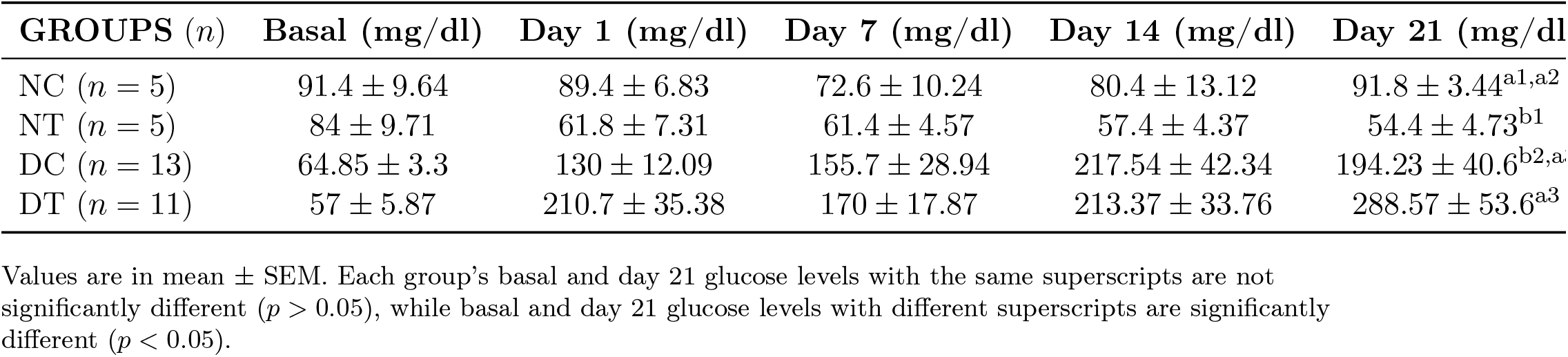
Fasting blood glucose of rats for the 21 days treatment period.

| GROUPS (*n*) | Basal (mg/dl) | Day 1 (mg/dl) | Day 7 (mg/dl) | Day 14 (mg/dl) | Day 21 (mg/dl) |
| --- | --- | --- | --- | --- | --- |
| NC (*n* = 5) | 91.4 ± 9.64 | 89.4 ± 6.83 | 72.6 ± 10.24 | 80.4 ± 13.12 | 91.8 ± 3.44<sup>a1,a2</sup> |
| NT (*n* = 5) | 84 ± 9.71 | 61.8 ± 7.31 | 61.4 ± 4.57 | 57.4 ± 4.37 | 54.4 ± 4.73<sup>b1</sup> |
| DC (*n* = 13) | 64.85 ± 3.3 | 130 ± 12.09 | 155.7 ± 28.94 | 217.54 ± 42.34 | 194.23 ± 40.6<sup>b2,a</sup> |
| DT (*n* = 11) | 57 ± 5.87 | 210.7 ± 35.38 | 170 ± 17.87 | 213.37 ± 33.76 | 288.57 ± 53.6<sup>a3</sup> |

Values are in mean ± SEM. Each group's basal and day 21 glucose levels with the same superscripts are not significantly different ( $p > 0.05$ ), while basal and day 21 glucose levels with different superscripts are significantly different ( $p < 0.05$ ).

### 3.3 Effects SOD and Catalase Activities

#### Effect on Pancreas SOD Activity

The effect of lemongrass extract on pancreas SOD activity is presented in Table 3.3. Relative to the control groups (NC and DC), lemongrass did not significantly decrease (*P >* 0.05) pancreas SOD activity in the treated groups (NT and DT).

**Table 3:** Enzyme activity for pancreas SOD in normal and diabetic rats (U/mg protein)

| GROUPS | TREATMENT | SOD Activity (U/mg) |
| --- | --- | --- |
| Normal Control | Water | 0.226 ± 0.064<sup>a1,a2</sup> |
| Normal Treated | Extract (0.4 g/kg b. wt.) | 0.177 ± 0.027<sup>a1</sup> |
| Diabetic Control | Streptozotocin (1ml/kg b. wt.) | 0.141 ± 0.034<sup>a2,a</sup> |
| Diabetic Treated | Streptozotocin + Extract | 0.204 ± 0.052<sup>a3</sup> |

Values are in mean ± SEM. Data with the same superscripts are not significantly different ( $p > 0.05$ ), while data with different superscripts are significantly different ( $p < 0.05$ ).

#### Effect on Pancreas Catalase Activity

The effect of lemongrass extract on pancreas catalase activity is presented in Table 3.4. Comparing results from the control groups (NC and DC) to that of the treated groups (NT and DT), it showed lemongrass did not significantly reduce (*P >* 0.05) pancreas catalase activity.

**Table 4:** Enzyme activity for pancreas catalase in normal and diabetic rats (U/mg protein)

| GROUPS | TREATMENT | Catalase Activity (U/mg) Mean ± SEM |
| --- | --- | --- |
| Normal Control | Water | 6.60 ± 1.05<sup>a1,a2</sup> |
| Normal Treated | Extract (0.4 g/kg b. wt.) | 6.56 ± 1.17<sup>a1</sup> |
| Diabetic Control | Streptozotocin (1ml/kg b. wt.) | 10.35 ± 6.46<sup>a2,a</sup> |
| Diabetic Treated | Streptozotocin + Extract | 7.55 ± 2.94<sup>a3</sup> |

Values are in mean ± SEM. Data with the same superscripts are not significantly different ( $p > 0.05$ ), data with different superscripts are significantly different ( $p < 0.05$ ).

## 4 Discussion

The study aimed to determine if 0.4 g/kg body weight of *Cymbopogon citratus* extract could reverse the diabetic effects induced by STZ (45 mg/kg body weight).

The findings confirm that STZ successfully induced diabetes, as both DC and DT rats showed significantly elevated FBG (above 200 mg/dl after day 14) and significantly lower weight gain compared to normal rats. A reason could be that the anti-diabetic components in lemongrass (citral, limonene and linalool), were unable to reverse the diabetic effect of STZ (Garba et al., 2020).

### Antidiabetic Effect

The extract successfully reduced blood glucose in normal rats (NT group) compared to the NC group (*P <* 0.05) but failed to significantly reduce blood glucose in the diabetic rats (DT group) compared to the DC group (*P >* 0.05). This suggests the anti-diabetic components of lemongrass, such as citral and limonene, were insufficient to reverse the *β*-cell destructive effects of STZ (Garba et al., 2020).

### Antioxidant Enzyme Activities (SOD and CAT)

The lack of statistically significant difference (*P >* 0.05) in SOD and CAT activities between all groups (NC, NT, DC, DT) suggests that the lemongrass extract had no statistically significant effect on the oxidative status of the pancreas in either normal or diabetic rats. This contrasts with some previous studies where STZ treatment alone caused elevated SOD levels (Thomas et al., 2014). The observation of similar SOD and CAT levels between the NC and NT groups implies the extract did not exert oxidative damage to the pancreas in normal rats.

In the DC rats, the observed reduced levels of pancreatic SOD (albeit insignificant) is hypothesized to be due to glucose auto-oxidation, which forms hydrogen peroxide that inactivates SOD, thus leading to its decreased activity (Fujita, 2009).

## 5 Conclusion

The results indicate that the aqueous extract of *Cymbopogon citratus* extract (400 mg/kg body weight) was ineffective in reversing STZ-induced diabetes in rats. While the extract showed a hypoglycemic effect in normal rats, it did not significantly increase body weight or decrease blood glucose in diabetic rats, nor did it cause oxidative damage to the pancreas.

While this study provides empirical data on the plant’s effect, most available studies are animal-based. Therefore, more empirical studies, particularly those evaluating the effect of *Cymbopogon citratus* on humans, are required to substantiate its use in therapeutics for diabetes management.

## Supporting information

Result data

---

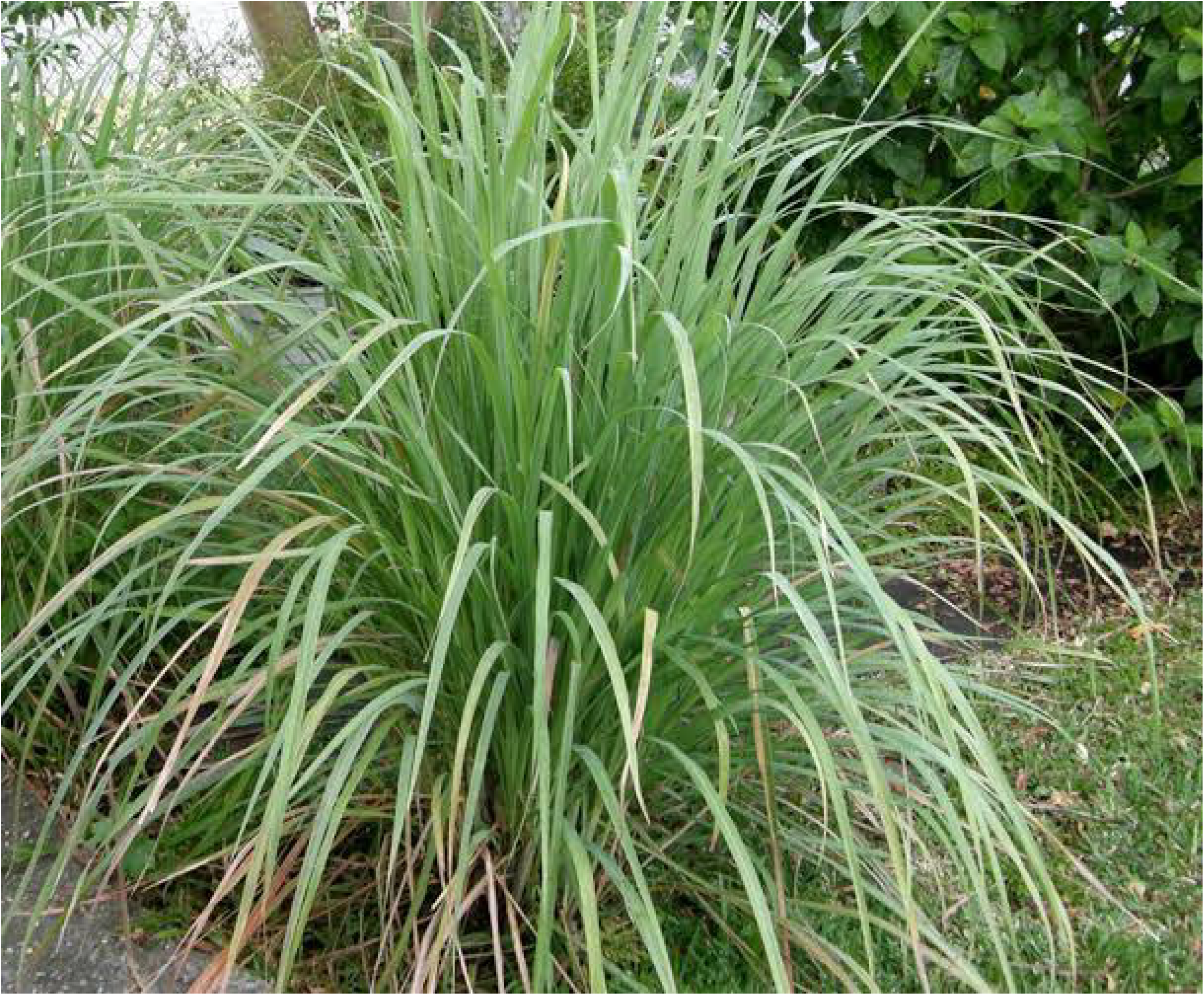

