## Supplementary material for "EFFECTS OF AQUEOUS EXTRACT OF *CYMBOPOGON CITRATUS* ON BLOOD GLUCOSE, BODY WEIGHT, AND PANCREATIC ANTIOXIDANT ACTIVITIES IN NORMAL AND STREPTOZOTOCIN-INDUCED DIABETIC RATS": Result data

### EFFECTS OF *CYMBOPOGON CITRATUS* ON NORMAL AND STREPTOZOTOCIN-INDUCED DIABETIC RATS BY DETERMINING BLOOD GLUCOSE, WEIGHT GAIN/LOSS AND PANCREATIC SUPEROXIDE DISMUTASE (SOD) AND CATALASE (CAT) ACTIVITIES

#### WEIGHT OF RATS

Table 1 showing weight of rats used for this study

**Normal control group**

| NAME | INITIAL WEIGHT (g) | FINAL WEIGHT (g) |
|---|---|---|
| H/B | 114.2 | 198 |
| T | 131.9 | 174 |
| B | 113.8 | 192 |
| H/T | 116.15 | 178 |
| H | 132.9 | 203 |

**Diabetic control group**

| NAMES | INITIAL WEIGHT (g) | FINAL WEIGHT (g) |
|---|---|---|
| | CAGE 1 | |
| H/T | 142.4 | 178 |
| H/B | 135.58 | 174 |
| A | 139.5 | 192 |
| A/RL | 146.65 | 136 |
| 2L | 147.11 | 163 |
| T/2L | 147 | 134 |
| H/2L | 150.4 | 185 |
| | CAGE 2 | |
| H/T | 134.58 | 170 |
| H | 140.41 | 134 |
| B | 146.62 | 144 |
| H/A | 139.13 | 123 |
| B/T | 133 | 167 |
| H/RL | 132.5 | 133 |

Normal treated group

| NAME | INITIAL WEIGHT (g) | FINAL WEIGHT (g) |
|---|---|---|
| H/B | 133.01 | 166 |
| H/RL | 128.61 | 203 |
| H/T | 122.11 | 198 |
| B | 128.28 | 174 |

| | | |
|---|---|---|
| A | 120.55 | 185 |

Diabetic treated group

| NAME | INITIAL WEIGHT (g) | FINAL WEIGHT (g) |
|---|---|---|
| | CAGE 1 | |
| A | 137.92 | 98 |
| B/RL | 155.16 | 144 |
| T/2L | 149.4 | 148 |
| 2L | 140.05 | 166 |
| H/B/RL | 140.1 | 185 |
| | CAGE 2 | |
| H/2L | 136.1 | 141 |
| H/A | 159 | 140 |
| H | 155 | 200 |
| RL | 144.7 | 179 |
| T | 158.3 | 199 |
| A | 162.9 | 132 |

GLUCOSE LEVELS OF RATS

Table showing the various glucose levels of rats used for the study

Normal control

| Name | Basal glucose (mg/dl) | Day 1 (mg/dl) | Day 7 (mg/dl) | Day 14 (mg/dl) | Day 21 (mg/dl) |
|---|---|---|---|---|---|
| H/B | 116 | 100 | 111 | 108 | 85 |
| T | 78 | 79 | 75 | 116 | 86 |
| B | 112 | 110 | 64 | 52 | 104 |
| H/T | 67 | 73 | 53 | 64 | 90 |
| H | 84 | 85 | 60 | 62 | 94 |

Normal treated

| Name | Basal glucose (mg/dl) | Day1 (mg/dl) | Day 7 (mg/dl) | Day 14 (mg/dl) | Day 21 (mg/dl) |
|---|---|---|---|---|---|
| H/B | 48 | 85 | 59 | 67 | 61 |
| H/RL | 83 | 39 | 59 | 46 | 39 |
| H/T | 87 | 63 | 50 | 51 | 52 |
| B | 100 | 63 | 78 | 55 | 67 |
| A | 102 | 59 | 61 | 68 | 53 |

Diabetic control

| Name | Basal glucose (mg/dl) | Day 1 (mg/dl) | Day 7 (mg/dl) | Day14 (mg/dl) | Day 21 (mg/dl) |
|---|---|---|---|---|---|
| | | Cage 1 | | | |
| H/T | 46 | 73 | 80 | 111 | 73 |
| H/B | 68 | 191 | 88 | 73 | 77 |
| A | 87 | 91 | 115 | 124 | 92 |
| A/RL | 78 | 148 | 105 | 358 | 134 |
| 2L | 67 | 157 | 111 | 346 | 122 |
| T/2L | 61 | 156 | 148 | 442 | 192 |
| H/2L | 61 | 132 | 80 | 96 | 107 |
| | | Cage 2 | | | |
| H/T | 46 | 76 | 58 | 92 | 88 |
| H | 62 | 151 | 401 | 192 | 324 |
| B | 56 | 92 | 248 | 438 | 535 |
| H/A | 74 | 211 | 251 | 398 | 370 |
| B/T | 62 | 101 | 68 | 54 | 91 |
| H/RL | 75 | 111 | 271 | 104 | 320 |

Diabetic treated

| Name | Basal glucose (mg/dl) | Day 1 (mg/dl) | Day 7 (mg/dl) | Day 14 (mg/dl) | Day 21 (mg/dl) |
|---|---|---|---|---|---|
| | | Cage 1 | | | |
| A | 45 | 464 | 160 | 128 | 315 |
| B/RL | 33 | 207 | 296 | 146 | 169 |
| T/2L | 51 | HI | 199 | 118 | 273 |

| | | | | | |
|---|---|---|---|---|---|
| 2L | 51 | 174 | 120 | 342 | 74 |
| H/B/RL | 52 | 156 | 124 | 125 | 151 |
| **Cage 2** | | | | | |
| H/2L | 37 | 172 | 147 | 112 | 487 |
| H/A | 64 | 187 | 213 | 403 | 433 |
| H | 59 | 83 | 135 | 322 | 520 |
| RL | 103 | 132 | 83 | 284 | HI |
| T | 77 | 348 | 173 | 96 | 175 |
| A | 55 | 184 | 220 | 271 | HI |

**CALCULATIONS SHOWING AMOUNTS OF STZ, CITRATE BUFFER, EXTRACT GIVEN TO EACH GROUP**

1. Normal control group:

The rats were given citrate buffer 1ml/kg body weight only. The volume given to each rat was calculated using the formula below:

1ml = 1000g

Xml = weight of rat

Therefore, Xml = $\frac{1\text{ml} \times \text{weight of rats}}{1000\text{g}}$ **...eq. 1**

*Table of normal control citrate buffer*

| Name | Buffer (ml) |
|---|---|
| H/B | 0.104 |
| T | 0.110 |
| B | 0.102 |
| H/T | 0.102 |
| H | 0.107 |

2. Diabetic control group:

The rats were given the STZ (45mg/kg) dissolved in 0.1M citrate buffer (1ml/kg) solution. The volume of the solution given to each rat was calculated using this formula below:

The amount of STZ dissolved in buffer

45mg = 1ml

Xmg = Zml

Therefore Zml = 0.022Xmg ..... **eq. 2.1**

Amount of STZ used given per rat

45mg = 1000g

Xmg = weight of rats

Therefore, $Xmg = \frac{45mg \times \text{weight of rats}}{1000g}$....**equ 2.2**

Where, X = amount of STZ in mg

Z = amount of citrate buffer in ml

*Table of diabetic control rats STZ*

| Name | STZ/citrate buffer solution (ml) |
|---|---|
| **Cage 1** | |
| H/T | 0.14 |
| H/B | 0.14 |
| A | 0.14 |
| A/RL | 0.15 |
| 2L | 0.15 |
| T/2L | 0.15 |
| H/2L | 0.14 |
| **Cage 2** | |
| H/T | 0.13 |
| H | 0.14 |
| B | 0.15 |
| H/A | 0.14 |
| B/T | 0.13 |
| H/RL | 0.13 |

3. Normal treated group:

**Buffer**

The rats were given the citrate buffer 1ml/kg refer to **equ 1:**

**Extract**

The rats were also given extract 400mg/kg body weight. The volume of extract given was calculated using the formula below:

The extract was dissolved in water (5% stock solution) 5g/100ml.

5g of extract = 100ml of water

400mg of extract = xml of water

Therefore 400mg = 8ml of water = 1000g rat

1000g = 8ml

Weight of rat = Eml

Therefore, Eml = $\frac{8\text{ml} \times \text{weight of rat}}{1000\text{g}}$...**eq. 3**

Where, E= amount of extract solution for each rat

**Values of each rat are represented below:**

| Name | Citrate buffer (ml) | Extract dose (ml) |
|---|---|---|
| H/B | 0.13 | 1.06 |
| H/RL | 0.13 | 1.03 |
| H/T | 0.12 | 0.98 |
| B | 0.13 | 1.03 |
| A | 0.12 | 0.96 |

4. Diabetic treated group:

These rats were given both the STZ solution, refer to **eq. 3** and the extract solution, refer to **eq. 2.1** and **eq. 2.2**

Value for each rat represented below:

| Name | Citrate buffer / STZ solution (ml) | Extract (ml) |
|---|---|---|
| **Cage 1** | | |
| A | 0.14 | 1.1 |
| B/RL | 0.15 | 1.2 |
| T/2L | 0.15 | 1.2 |
| 2L | 0.14 | 1.1 |
| H/B/RL | 0.16 | 1.2 |
| **Cage 2** | | |
| H/2L | 0.14 | 1.1 |
| H/A | 0.16 | 1.3 |
| H | 0.16 | 1.2 |
| RL | 0.15 | 1.2 |
| T | 0.16 | 1.3 |
| A | 0.16 | 1.3 |

#### 2.1 BIOCHEMICAL ASSAYS

#### A. SOD ASSAY

The activity of catalase in each sample is calculated as:

$$\% \text{ Inhibition} = \frac{O.D_{test} - O.D_{reference}}{O.D_{test}} \times \frac{100}{1}$$

$$\text{Enzyme Activity (units/mg protein)} = \frac{\% \text{ inhibition}}{50 \times Y}$$

Where Y = mg of protein in the volume of sample.

$O.D_{reference} = 0.007$

A unit of SOD activity was taken as the amount of SOD required to cause 50 % inhibition of the auto-oxidation of adrenaline to adrenochrome per minute.

**Solving for H/B (NC)**

$$\% \text{ Inhibition} = \frac{(0.037 - 0.007) \times 100}{0.037}$$

% Inhibition = 81.08%

Enzyme activity = 81.08 ÷ (50 × 3.585)

Enzyme activity = 0.452 U/mg protein.

***The same process was applied to every specimen.**

*Table 2a showing SOD absorbance*

| | | | | | | |
|---|---|---|---|---|---|---|
| | SOD RESULTS (O. D) | | | | | |
| | ABSORBANCE @ 420nm | | | | | |
| **Normal Control** | **1mins** | **1mins** | **2mins** | **2mims** | **3mins** | **3mins** |
| H/B | 1.615 | 1.611 | 1.646 | 1.646 | 1.65 | 1.65 |
| T | 1.575 | 1.571 | 1.65 | 1.65 | 1.65 | 1.656 |
| B | 1.476 | 1.472 | 1.63 | 1.636 | 1.646 | 1.65 |
| A/T | 1.465 | 1.461 | 1.645 | 1.645 | 1.65 | 1.65 |
| H | 1.574 | 1.57 | 1.645 | 1.651 | 1.65 | 1.654 |
| **Normal Treated** | | | | | | |
| H/B | **1.601** | **1.605** | 1.633 | 1.651 | 1.647 | 1.657 |
| H/RL | 1.49 | 1.494 | 1.648 | 1.648 | 1.651 | 1.655 |
| H/T | 1.507 | 1.511 | 1.65 | 1.648 | 1.653 | 1.653 |
| B | 1.497 | 1.5 | 1.646 | 1.646 | 1.64 | 1.658 |
| A | 1.558 | 1.562 | 1.64 | 1.65 | 1.65 | 1.65 |
| **Diabetic Control Cage 1** | | | | | | |
| H/T | **1.603** | **1.615** | 1.621 | 1.627 | 1.644 | 1.644 |
| H/B | 1.614 | 1.626 | 1.65 | 1.65 | 1.657 | 1.651 |
| A | 1.517 | 1.529 | 1.65 | 1.65 | 1.654 | 1.654 |
| A/RL | 1.53 | 1.542 | 1.648 | 1.648 | 1.656 | 1.65 |
| 2L | 1.541 | 1.541 | 1.636 | 1.636 | 1.65 | 1.658 |
| T/2L | 1.609 | 1.62 | 1.64 | 1.654 | 1.66 | 1.656 |
| H/2L | 0.982 | 0.964 | 1.387 | 1.387 | 1.568 | 1.568 |
| **Diabetic control cage 2** | | | | | | |
| H/T | **1.62** | **1.61** | 1.654 | 1.64 | 1.65 | 1.65 |
| H | 1.647 | 1.657 | 1.633 | 1.633 | 1.688 | 1.68 |
| B | 1.518 | 1.528 | 1.63 | 1.656 | 1.653 | 1.651 |
| H/A | 1.362 | 1.372 | 1.64 | 1.64 | 1.65 | 1.654 |
| B/T | 1.604 | 1.614 | 1.641 | 1.657 | 1.646 | 1.66 |
| H/RL | 1.25 | 1.26 | 1.492 | 1.492 | 1.618 | 1.64 |
| **Diabetic Treated Cage1** | | | | | | |
| A | **1.54** | **1.53** | 1.653 | 1.653 | 1.656 | 1.656 |
| B/RL | 1.513 | 1.503 | 1.654 | 1.654 | 1.65 | 1.66 |
| T/2L | 1.537 | 1.527 | 1.646 | 1.646 | 1.644 | 1.66 |
| 2L | 1.625 | 1.615 | 1.648 | 1.64 | 1.66 | 1.65 |
| H/B/RL | 1.575 | 1.565 | 1.649 | 1.649 | 1.653 | 1.653 |
| **Diabetic Treated Cage2** | | | | | | |
| H/2L | **1.55** | **1.558** | 1.648 | 1.648 | 1.656 | 1.653 |
| H/A | 1.456 | 1.44 | 1.65 | 1.65 | 1.54 | 1.54 |
| H | 1.631 | 1.631 | 1.652 | 1.648 | 1.653 | 1.653 |
| RL | 1.56 | 1.557 | 1.65 | 1.65 | 1.65 | 1.656 |
| T | 1.591 | 1.595 | 1.65 | 1.656 | 1.65 | 1.656 |
| A | 1.493 | 1.493 | 1.64 | 1.64 | 1.654 | 1.65 |

*Table 2b showing SOD concentration and mean absorbance*

MEAN RESULTS

| Normal Control | 1mins | 2mins | 3mins | Diff. | % Inhibition | U/mg protein |
|---|---|---|---|---|---|---|
| H/B | 1.613 | 1.646 | 1.65 | 0.037 | 81.08 | 0.452 |
| T | 1.573 | 1.65 | 1.653 | 0.08 | 96.11 | 0.117 |
| B | 1.474 | 1.633 | 1.648 | 0.174 | 95.98 | 0.170 |
| A/T | 1.463 | 1.645 | 1.65 | 0.187 | 96.26 | 0.11 |
| H | 1.572 | 1.648 | 1.652 | 0.08 | 91.25 | 0.28 |
| **MEAN** | **1.539** | **1.644** | **1.651** | **0.112** | 92.14 | 0.226 |
| ± SEM | 0.030 | 0.003 | 0.002 | 0.001 | 2.92 | 0.064 |
| Normal Treated | | | | | | |
| H/B | 1.603 | 1.642 | 1.652 | 0.049 | 85.71 | 0.246 |
| H/RL | 1.492 | 1.648 | 1.653 | 0.161 | 95.65 | 0.096 |
| H/T | 1.509 | 1.649 | 1.653 | 0.144 | 95.14 | 0.144 |
| B | 1.499 | 1.646 | 1.649 | 0.150 | 95.33 | 0.225 |
| A | 1.560 | 1.635 | 1.650 | 0.090 | 93 | 0.173 |
| **MEAN** | **1.533** | **1.644** | **1.651** | **0.119** | 92.97 | 0.177 |
| **± SEM** | **0.050** | **0.002** | **0.001** | **0.021** | 1.873 | 0.027 |
| Diabetic Control Cage 1 | | | | | | |
| H/T | 1.609 | 1.624 | 1.644 | 0.035 | 80 | 0.102 |
| H/B | 1.620 | 1.650 | 1.654 | 0.034 | 79.41 | 0.088 |
| A | 1.523 | 1.650 | 1.654 | 0.131 | 94.66 | 0.096 |
| A/RL | 1.536 | 1.648 | 1.653 | 0.117 | 94.02 | 0.119 |
| 2L | 1.547 | 1.636 | 1.654 | 0.107 | 93.46 | 0.198 |
| T/2L | 1.615 | 1.647 | 1.658 | 0.043 | 83.72 | 0.202 |
| H/2L | 0.958 | 1.387 | 1.568 | 0.610 | 98.85 | 0.317 |
| **MEAN** | **1.487** | **1.606** | **1.641** | **0.154** | 89.16 | 0.160 |
| **± SEM** | **0.090** | **0.036** | **0.012** | **0.078** | 2.987 | 0.032 |
| Diabetic control cage 2 | | | | | | |
| H/T | 1.615 | 1.647 | 1.650 | 0.035 | 80 | 0.094 |
| H | 1.652 | 1.633 | 1.684 | 0.032 | 78.13 | 0.01 |
| B | 1.523 | 1.643 | 1.652 | 0.129 | 94.57 | 0.067 |
| H/A | 1.367 | 1.640 | 1.652 | 0.285 | 97.54 | 0.199 |
| B/T | 1.609 | 1.649 | 1.653 | 0.044 | 84.09 | 0.117 |
| H/RL | 1.255 | 1.492 | 1.629 | 0.374 | 98.13 | 0.242 |
| **MEAN** | **1.504** | **1.617** | **1.653** | **0.150** | 88.74 | 0.122 |
| **± SEM** | **0.065** | **0.025** | **0.007** | **0.060** | 3.7 | 0.035 |
| Diabetic Treated Cage1 | | | | | | |
| A | 1.535 | 1.653 | 1.656 | 0.121 | 94.21 | 0.212 |
| B/RL | 1.508 | 1.654 | 1.655 | 0.147 | 95.24 | 0.404 |
| T/2L | 1.532 | 1.646 | 1.652 | 0.120 | 94.17 | 0.293 |
| 2L | 1.620 | 1.644 | 1.655 | 0.035 | 80 | 0.067 |
| H/B/RL | 1.570 | 1.649 | 1.653 | 0.083 | 91.57 | 0.118 |
| **MEAN** | **1.553** | **1.649** | **1.654** | **0.101** | 91.04 | 0.219 |
| **± SEM** | **0.019** | **0.002** | **0.001** | **0.019** | 2.825 | 0.06 |
| Diabetic Treated Cage2 | | | | | | |
| H/2L | 1.554 | 1.648 | 1.653 | 0.099 | 92.93 | 0.064 |
| H/A | 1.448 | 1.650 | 1.540 | 0.092 | 92.39 | 0.251 |
| H | 1.631 | 1.650 | 1.653 | 0.022 | 68.18 | 0.157 |
| RL | 1.559 | 1.650 | 1.653 | 0.094 | 92.55 | 0.192 |
| T | 1.593 | 1.653 | 1.655 | 0.062 | 88.71 | 0.108 |
| A | 1.493 | 1.640 | 1.652 | 0.159 | 95.60 | 0.362 |
| **MEAN** | **1.546** | **1.649** | **1.634** | **0.088** | 88.39 | 0.189 |
| **± SEM** | **0.027** | **0.002** | **0.019** | **0.018** | 4.141 | 0.044 |

#### B. CATALASE ASSAY

The activity of catalase in each sample is calculated thus:

$$\frac{O.D/min \times Vt \times 1000}{M \times V \times L \times Y}$$

where,

O.D = Absorbance of sample test at 480 nm

Vt = Total volume of the reaction mixture = 13.5 mL

M = Molar extinction coefficient of $H_2O_2$ = $43.6M^{-1}\ cm^{-1}$

L = Light path = 1.0 cm

V = Volume of sample homogenate used = 0.5 mL

Y = mg of protein in tissue used

**Solving for H/B (NC)**

$$\text{Activity} = \frac{0.037 \times 13.5 \times 1000}{43.6 \times 0.5 \times 1 \text{ x } 17.74}$$

Activity = 6.39 U/mg protein

***The same process was applied to every specimen.**

*Table 2c showing Catalase absorbance*

| | | | | | | |
|---|---|---|---|---|---|---|
| | | | CATALASE O.D RESULT | | | |
| | | | ABSORBANCE @ 480nm | | | |
| **Normal Control** | **0.5mins** | **0.5mins** | **1.5mins** | **1.5mins** | **2.5mins** | **2.5mins** |
| H/B | 1.6 | 1.626 | 1.64 | 1.652 | 1.652 | 1.648 |
| T | 1.55 | 1.596 | 1.65 | 1.65 | 1.655 | 1.651 |
| B | 1.483 | 1.465 | 1.631 | 1.634 | 1.65 | 1.646 |
| A/T | 1.463 | 1.462 | 1.665 | 1.629 | 1.652 | 1.648 |
| H | 1.591 | 1.552 | 1.641 | 1.655 | 1.654 | 1.65 |
| **Normal Treated** | | | | | | |
| H/B | 1.601 | 1.604 | 1.64 | 1.643 | 1.649 | 1.655 |
| H/RL | 1.502 | 1.482 | 1.649 | 1.646 | 1.65 | 1.656 |
| H/T | 1.5 | 1.517 | 1.65 | 1.657 | 1.65 | 1.656 |
| B | 1.505 | 1.494 | 1.646 | 1.646 | 1.647 | 1.652 |
| A | 1.56 | 1.56 | 1.63 | 1.64 | 1.65 | 1.656 |
| **Diabetic Control Cage 1** | | | | | | |
| H/T | 1.615 | 1.603 | 1.62 | 1.628 | 1.648 | 1.64 |
| H/B | 1.62 | 1.62 | 1.65 | 1.65 | 1.657 | 1.65 |
| A | 1.513 | 1.533 | 1.65 | 1.65 | 1.658 | 1.65 |
| A/RL | 1.536 | 1.535 | 1.652 | 1.65 | 1.657 | 1.649 |
| 2L | 1.55 | 1.544 | 1.641 | 1.631 | 1.658 | 1.65 |
| T/2L | 1.61 | 1.62 | 1.651 | 1.644 | 1.661 | 1.654 |
| H/2L | 0.965 | 0.951 | 1.4 | 1.373 | 1.562 | 1.564 |
| **Diabetic Control cage 2** | | | | | | |
| H/T | 1.61 | 1.62 | 1.65 | 1.644 | 1.65 | 1.65 |
| H | 1.65 | 1.654 | 1.645 | 1.62 | 1.682 | 1.684 |
| B | 1.546 | 1.5 | 1.636 | 1.65 | 1.65 | 1.653 |
| H/A | 1.368 | 1.365 | 1.64 | 1.64 | 1.65 | 1.654 |
| B/T | 1.61 | 1.607 | 1.65 | 1.647 | 1.651 | 1.655 |
| H/RL | 1.25 | 1.26 | 1.507 | 1.477 | 1.627 | 1.631 |
| **Diabetic Treated Cage1** | | | | | | |
| A | 1.53 | 1.54 | 1.65 | 1.655 | 1.659 | 1.653 |
| B/RL | 1.513 | 1.502 | 1.658 | 1.65 | 1.658 | 1.652 |
| T/2L | 1.551 | 1.513 | 1.642 | 1.65 | 1.655 | 1.649 |
| 2L | 1.62 | 1.62 | 1.655 | 1.633 | 1.658 | 1.652 |
| H/B/RL | 1.578 | 1.562 | 1.65 | 1.648 | 1.656 | 1.65 |
| **Diabetic Treated Cage2** | | | | | | |
| H/2L | 1.558 | 1.55 | 1.65 | 1.645 | 1.653 | 1.653 |
| H/A | 1.462 | 1.434 | 1.643 | 1.657 | 1.54 | 1.54 |
| H | 1.63 | 1.632 | 1.65 | 1.65 | 1.653 | 1.653 |
| RL | 1.567 | 1.551 | 1.65 | 1.65 | 1.652 | 1.653 |
| T | 1.595 | 1.59 | 1.65 | 1.656 | 1.65 | 1.66 |
| A | 1.48 | 1.506 | 1.647 | 1.633 | 1.654 | 1.65 |

*Table 2d showing Catalase concentrations and absorbance*

CAT. MEAN RESULTS

| Normal Control | 0.5mins | 1.5mins | 2.5mins | Diff. | U/mg protein |
|---|---|---|---|---|---|
| H/B | 1.613 | 1.646 | 1.65 | 0.037 | 6.39 |
| T | 1.573 | 1.65 | 1.653 | 0.08 | 3.02 |
| B | 1.474 | 1.633 | 1.648 | 0.174 | 9.52 |
| A/T | 1.463 | 1.645 | 1.65 | 0.187 | 6.60 |
| H | 1.572 | 1.648 | 1.652 | 0.08 | 7.50 |
| **MEAN** | **1.539** | **1.644** | **1.651** | **0.112** | **6.60** |
| **± SEM** | **0.030** | **0.003** | **0.002** | **0.001** | **1.05** |
| Normal Treated | | | | | |
| H/B | 1.603 | 1.642 | 1.652 | 0.049 | 4.35 |
| H/RL | 1.492 | 1.648 | 1.653 | 0.161 | 4.98 |
| H/T | 1.509 | 1.649 | 1.653 | 0.144 | 6.75 |
| B | 1.499 | 1.646 | 1.649 | 0.15 | 10.94 |
| A | 1.56 | 1.635 | 1.65 | 0.09 | 5.76 |
| **MEAN** | **1.533** | **1.644** | **1.651** | **0.119** | **6.56** |
| **± SEM** | **0.050** | **0.002** | **0.001** | **0.021** | **1.17** |
| Diabetic Control Cage 1 | | | | | |
| H/T | 1.609 | 1.624 | 1.644 | 0.035 | 1.38 |
| H/B | 1.620 | 1.650 | 1.654 | 0.034 | 1.16 |
| A | 1.523 | 1.650 | 1.654 | 0.131 | 4.13 |
| A/RL | 1.536 | 1.648 | 1.653 | 0.117 | 4.57 |
| 2L | 1.547 | 1.636 | 1.654 | 0.107 | 7.02 |
| T/2L | 1.615 | 1.647 | 1.658 | 0.043 | 3.21 |
| H/2L | 0.958 | 1.387 | 1.568 | 0.610 | 60.66 |
| **MEAN** | **1.487** | **1.606** | **1.641** | **0.154** | **11.73** |
| **± SEM** | **0.090** | **0.036** | **0.012** | **0.078** | **8.19** |
| Diabetic Control cage 2 | | | | | |
| H/T | 1.615 | 1.647 | 1.650 | 0.035 | 1.28 |
| H | 1.652 | 1.633 | 1.684 | 0.032 | 1.27 |
| B | 1.523 | 1.643 | 1.652 | 0.129 | 2.82 |
| H/A | 1.367 | 1.640 | 1.652 | 0.285 | 17.99 |
| B/T | 1.609 | 1.649 | 1.653 | 0.044 | 1.90 |
| H/RL | 1.255 | 1.492 | 1.629 | 0.374 | 28.54 |
| **MEAN** | **1.504** | **1.617** | **1.653** | **0.150** | **8.97** |
| **± SEM** | **0.065** | **0.025** | **0.007** | **0.060** | **4,73** |
| Diabetic Treated Cage1 | | | | | |
| A | 1.535 | 1.653 | 1.656 | 0.121 | 8.45 |
| B/RL | 1.508 | 1.654 | 1.655 | 0.147 | 19.30 |
| T/2L | 1.532 | 1.646 | 1.652 | 0.120 | 11.58 |
| 2L | 1.620 | 1.644 | 1.655 | 0.035 | 0.90 |
| H/B/RL | 1.570 | 1.649 | 1.653 | 0.083 | 3.32 |
| **MEAN** | **1.553** | **1.649** | **1.654** | **0.101** | **8.71** |
| **± SEM** | **0.019** | **0.002** | **0.001** | **0.019** | **3.24** |
| Diabetic Treated Cage2 | | | | | |
| H/2L | 1.554 | 1.648 | 1.653 | 0.099 | 2.10 |
| H/A | 1.448 | 1.650 | 1.540 | 0.092 | 7.74 |
| H | 1.631 | 1.650 | 1.653 | 0.022 | 1.57 |
| RL | 1.559 | 1.650 | 1.653 | 0.094 | 6.05 |
| T | 1.593 | 1.653 | 1.655 | 0.062 | 2.34 |
| A | 1.493 | 1.640 | 1.652 | 0.159 | 18.63 |
| **MEAN** | **1.546** | **1.649** | **1.634** | **0.088** | **6.40** |
| **± SEM** | **0.027** | **0.002** | **0.019** | **0.018** | **2.64** |

### STATISTICAL ANALYSIS

Using ANOVA, the P−value for the **NC−NT** set was calculated as;

Get External Data | Connections | Sort & Filter

Q3 | $f_x$ | 6.390447

| | A | B | C | D | E | F | G |
|---|---|---|---|---|---|---|---|
| 1 | Anova: Single Factor | | | | | | |
| 2 | | | | | | | |
| 3 | SUMMARY | | | | | | |
| 4 | *Groups* | *Count* | *Sum* | *Average* | *Variance* | | |
| 5 | Column 1 | 5 | 36.79571 | 7.359142 | 1.629039 | | |
| 6 | Column 2 | 5 | 32.77661 | 6.555321 | 6.809538 | | |
| 7 | | | | | | | |
| 8 | | | | | | | |
| 9 | ANOVA | | | | | | |
| 10 | *Source of Variation* | *SS* | *df* | *MS* | *F* | *P-value* | *F crit* |
| 11 | Between Groups | 1.615322 | 1 | 1.615322 | 0.382842 | 0.553292 | 5.317655 |
| 12 | Within Groups | 33.7543 | 8 | 4.219288 | | | |
| 13 | | | | | | | |
| 14 | Total | 35.36963 | 9 | | | | |
| 15 | | | | | | | |

***The same process was applied to other set of groups.**

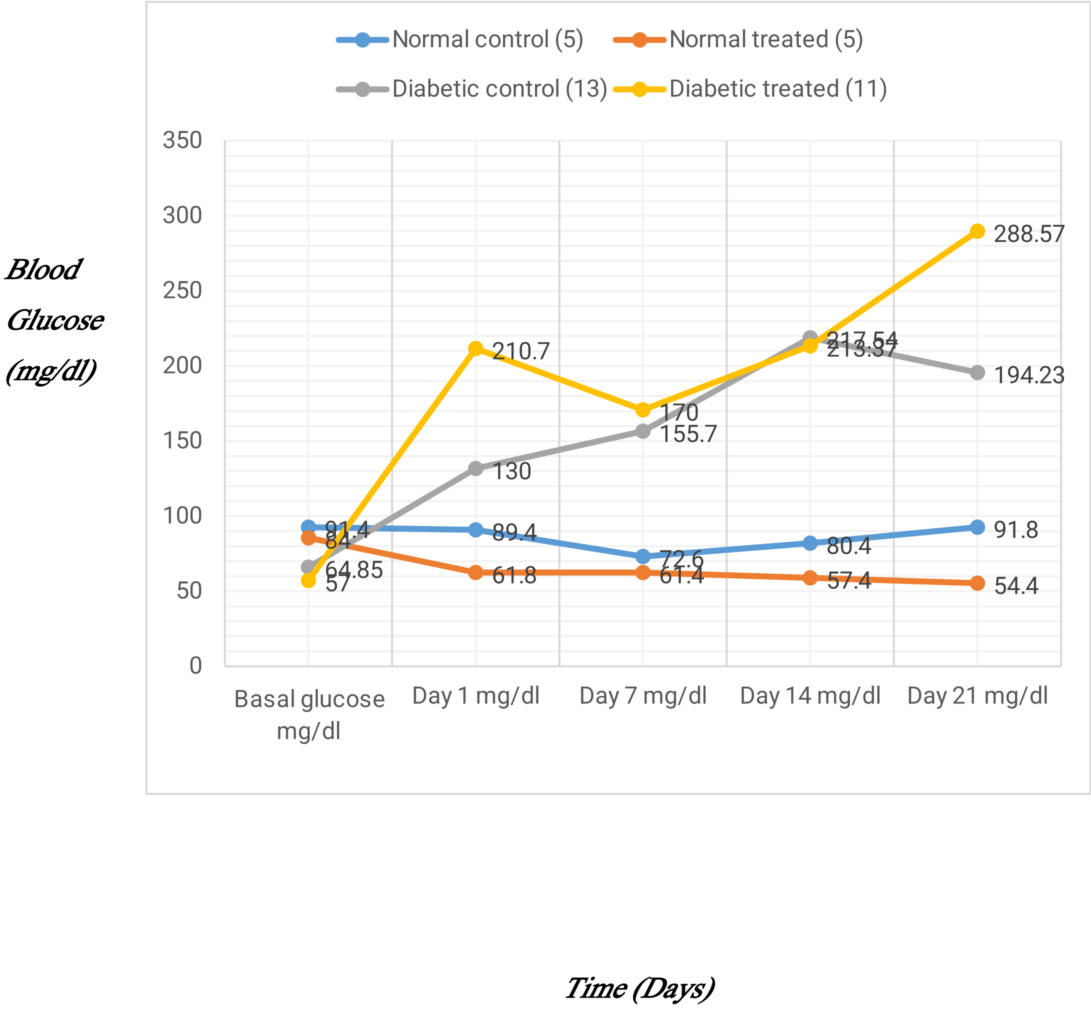


Chart 1: Glucose levels during 21 days treatment.

Weig-

ht (g)

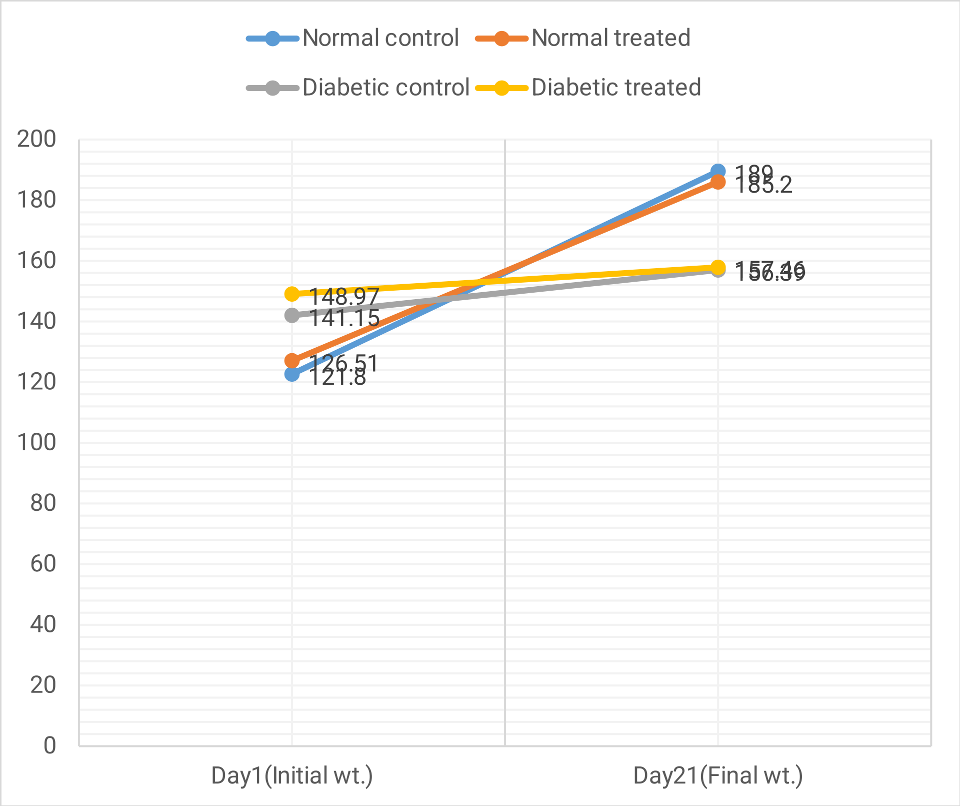


Time (Days)

**Chart 2: Specimen weights before and after 21 days.**
